# Neural Tracking of Audiovisual Effects in Noise Using Deep Neural Network-Generated Virtual Humans

**DOI:** 10.1101/2025.06.02.656280

**Authors:** John Kyle Cooper, Jonas Vanthornhout, Astrid van Wieringen, Tom Francart

## Abstract

This study investigates the effectiveness of Deep Neural Network (DNN)-generated virtual humans in enhancing audiovisual speech perception in noisy environments, with a focus on using neural measures to quantify these effects. Lip movements are essential for speech comprehension, especially when auditory cues are degraded by noise. Traditional recording methods produce high-quality audiovisual materials but are resource intensive. This research explores the use of DNN avatars as a promising alternative, utilizing a commercially available tool to create realistic virtual humans. The study included both simple sentences and a short story to improve ecological validity.

Eleven young, normal-hearing participants proficient in Flemish-Dutch listened to semantically meaningful sentences and a short story with various speaker types: a female FACS avatar, male and female DNN avatars, and a video of a human male speaker. The study included behavioral measures which consisted of an adaptive recall procedure and an adaptive rate procedure and electrophysiological measures consisting of neural tracking.

Findings in the adaptive recall procedure showed consistent audiovisual benefits, with the human speaker offering the greatest benefit (−4.75 dB SNR), followed by the DNN avatar (−4.00 dB SNR) and the FACS avatar (−1.55 dB SNR). Additionally in the adaptive rate procedure, the DNN avatar improved speech intelligibility, with average SRTs enhancing from −7.17 dB SNR (audio-only) to −9.02 dB SNR (audiovisual). The results from the neural tracking procedure indicated that most participants experienced audiovisual benefits, particularly in the −9 dB SNR range, revealing that audiovisual cues provided by DNN avatars can enhance speech perception, validating these avatars as effective tools for studying audiovisual effects when using both behavioral measures and electrophysiological measures.

## Introduction

Lip movements play a crucial role in speech comprehension, particularly in challenging acoustic environments, by providing visual cues that enhance auditory perception (Sumby & Pollack, 1954; Grant & Seitz, 2000; Peelle & Sommers, 2015). For individuals with hearing loss, there is a need for ecologically valid measures of speech understanding that reflect real-world communication conditions, including audiovisual speech cues (Pichora-Fuller et al., 2016). While behavioral methods can measure the benefits of lip movements using live speakers (Sumby & Pollack, 1954) or video recordings (Grant & Seitz, 2000; Ross et al., 2007; Llorach et al., 2022; Le Rhun et al., 2024), the production of high-quality audiovisual speech materials presents significant challenges, both in terms of time and resource investment (Akeroyd et al., 2015). Moreover, existing auditory-only speech materials cannot be added to visual speech material, which requires the creation of entirely new audiovisual corpora.

As a solution researchers have increasingly turned to synthetic audiovisual generation techniques, with virtual humans or avatars emerging as a promising solution for producing new audiovisual speech materials from existing auditory-only speech materials (Mattheyses & Verhelst, 2015). For generating these virtual humans, we present two primary approaches. The first method employs 3D modeling techniques to create realistic human representations, with visual cues controlled through various methods: visual articulators (Massaro & Cohen, 1990; Salvi et al., 2009), anatomy-based deformation systems (Sifakis et al., 2005), or 3D facial mapping using motion capture techniques (Williams, 1990; Fagel et al., 2007). For this study, we categorize these 3D modeling techniques as Facial Action Coding Systems (FACS) (Ekman & Friesen, 1976). As a broad concept, FACS is a system that simulates facial musculature in 3D modeled virtual humans to create more accurate lip movements. The second approach focuses on transforming acoustic speech into synthetic visual cues paired with a virtual human, a methodology extensively explored through evolving statistical machine learning techniques (Salvi et al., 2009; Massaro et al., 1999; Beskow et al., 2002, 2004). Building from these statistical machine learning techniques, recent advancements have increasingly utilized deep neural network (DNN) approaches to enhance the generation of synthetic visual cues (Eskimez et al., 2018; Vougioukas et al., 2020; Eskimez et al., 2020, 2022).

In the domain of FACS virtual humans, notable examples include MASSY (Fagel and Clemens, 2004), which investigated audiovisual cue perception in both normal-hearing and hearing-impaired individuals, and AVATAR (Devesse et al., 2018; Devesse, van Wieringen, et al., 2020a, 2020b; Devesse, Wouters, et al., 2020; Porto et al., 2025), a validated paradigm using semantically predictable sentences recited by virtual avatars in realistic environments (e.g., a restaurant scenario). The AVATAR paradigm has been comprehensively evaluated for both normal-hearing and hearing-impaired individuals. These studies show that FACS virtual humans improve SRTs but less than a video recording of a real human speaker.

Regarding virtual humans generated through DNN techniques, researchers have investigated audiovisual benefit from realistic virtual humans that recite existing speech materials (Shan et al., 2022; Varano et al., 2022; Yu et al., 2024). Their research demonstrated significant audiovisual benefits for listeners, highlighting the potential of synthetic visual cue generation. However, like FACS virtual humans, these studies found that these DNN virtual humans do not provide as much audiovisual benefit as a real human speaker.

In our study, we initially utilize the AVATAR paradigm; however, the avatars in this paradigm are only programmed to recite specific speech materials. To enhance the realism of this paradigm, we experimented with transitioning from a FACS-based avatar to a DNN-based avatar. The key advantage of a DNN-based avatar is its ability to process any speech material as input, generating realistic audiovisual outputs. To further improve listener engagement and make our study more ecological, we also moved beyond simple sentences and incorporated a short story into our study. We employed a commercially available tool, HeyGen (*HeyGen - AI Spokesperson Video Creator*, n.d.), to create realistic avatars, which were then tested. The commercial model utilized in this study operates on a proprietary architecture; however, prior studies have demonstrated that a specific type of deep neural network, known as generative adversarial networks (GANs), yields substantial audiovisual benefits (Shan et al., 2022; Varano et al., 2022; Yu et al., 2024).

While recent studies have investigated behavioral measures of audiovisual benefit of virtual humans, more objective measures of audiovisual benefit using neural responses measured with magneto/electroencephalography (M/EEG) have not been extensively studied. The primary reason for pursuing these more objective measures is that behavioral measures rely on input from the participant. More specifically, with more naturalistic speech material behavioral measures typically depend on an imprecise and subjective ratings. In contrast, more objective measures using neural responses do not require any input from the participant, which has many advantages such as enhancing participant engagement and enabling work with participant populations who may be unable to provide input.

One of these more objective measures of neural responses associated with a stimulus is called neural tracking, which has been shown to reflect speech intelligibility in both quiet and noisy environments (Vanthornhout et al., 2018; Lesenfants et al., 2019; Gillis et al., 2021). Neural tracking methods allow researchers to investigate neural responses using continuous stimuli, which is an advantage over traditional event related potential (ERP) methods (Lalor et al., 2009). Additional studies have found that observing mouth movements while listening to speech enhances neural tracking, especially when the movements align with the speech, compared to conditions with mismatched or absent mouth movements (Zion Golumbic et al., 2013; Crosse et al., 2015; O’Sullivan et al., 2019, 2021; Ahmed et al., 2023). Further studies have shown that neural tracking to audiovisual speech can predict behavioral outcomes such as multisensory integration benefit (Crosse et al., 2016) and the impact of visual cue removal on comprehension and perceived difficulty (Reisinger et al., 2025) and may correlate with speech reception thresholds (Cooper et al., 2025).

Other recent research includes the study of audiovisual illusory effects by using intracranial EEG recordings, naturalistic speech stimuli, and FACS virtual humans to provide a more ecologically valid way to investigate audiovisual speech processing (Thézé et al., 2020; Mégevand et al., 2025).

Previous research has demonstrated that for normal hearing participants, avatars provide an audiovisual benefit that can be measured both behaviorally and through neural tracking (Cooper et al., 2025). Therefore, our goal with the current study was to first compare the DNN-based avatars with our FACS-based avatar and a video of a human speaker to assess any improvements in speech intelligibility among the study participants. Based on previous research we hypothesized that the FACS and DNN-based avatars would provide similar audiovisual benefit to listener and the video of a human speaker would provide the most audiovisual benefit to the listener. Furthermore, we then validated whether neural tracking can demonstrate an audiovisual benefit when participants watch the DNN-based avatar recite a short story. Based on previous research, we hypothesized that these DNN-based avatars would yield comparable benefits to previous studies in both behavioral and electrophysiological measures.

## Methods

### Participants

For our study we recruited 11 young (ages 18-25) participants (5 female, 6 male) with normal hearing, language processing, and cognitive function. Normal hearing was defined as having hearing thresholds of 20 dB HL or below across the frequency range of 125 Hz to 8000 Hz, verified for each participant using a MADSEN Itera II. All participants were proficient in Flemish-Dutch and completed both behavioral and electrophysiological tests, which were administered sequentially in that order. The medical ethics committee of the UZ Leuven/KU Leuven approved this experiment and every participant signed an informed consent form before their participation (S65640).

### Speech Stimuli

For our speech stimuli, we utilized LIST sentences (van Wieringen & Wouters, 2008; Jansen et al., 2014) and a short story. The LIST sentences have been previously employed to compare behavioral (Devesse, van Wieringen, et al., 2020a) and neural tracking results (Cooper et al., 2025) within the AVATAR paradigm. In our study, we used two versions of the LIST sentences—one female and one male—ensuring that there was no overlap in the sentences selected from each version. The short story, a children’s narrative (*Milan • deBuren*, n.d.) of 13 minutes, has also been used in previous neural tracking studies (Vanthornhout et al., 2018; De Clercq et al., 2023). We truncated silences that exceeded 400 ms (Ding & Simon, 2012; Maddox & Lee, 2018) and divided the story into 1-minute segments. To maximize the number of segments, we did not adjust the segments to include the endings of sentences, resulting in segments that may not have corresponded to complete sentences.

### Speakers (avatars and humans)

In this study, we compare the audiovisual benefit provided by a human speaker to that of virtual human speakers. For the virtual humans, we utilized the existing AVATAR paradigm, incorporating two different types of avatars. The first is the female avatar created using the AVATAR paradigm, which uses a FACS 3D-modeled human. The second avatar type was generated using a commercial avatar creation tool that employs deep neural networks (DNNs), to create virtual humans with realistic lip movements for any speech material (*HeyGen - AI Spokesperson Video Creator*, n.d.). These DNN avatars exhibit a full range of body movements driven by the input speech, including head, arm, and hand movements. Using this tool, we developed both a female and a male avatar to recite the LIST sentences. In addition, we included a video recording of a human male speaker reciting the LIST sentences. Visual representations of the different speaker types are displayed in Figure 1B.

**Figure 1.**
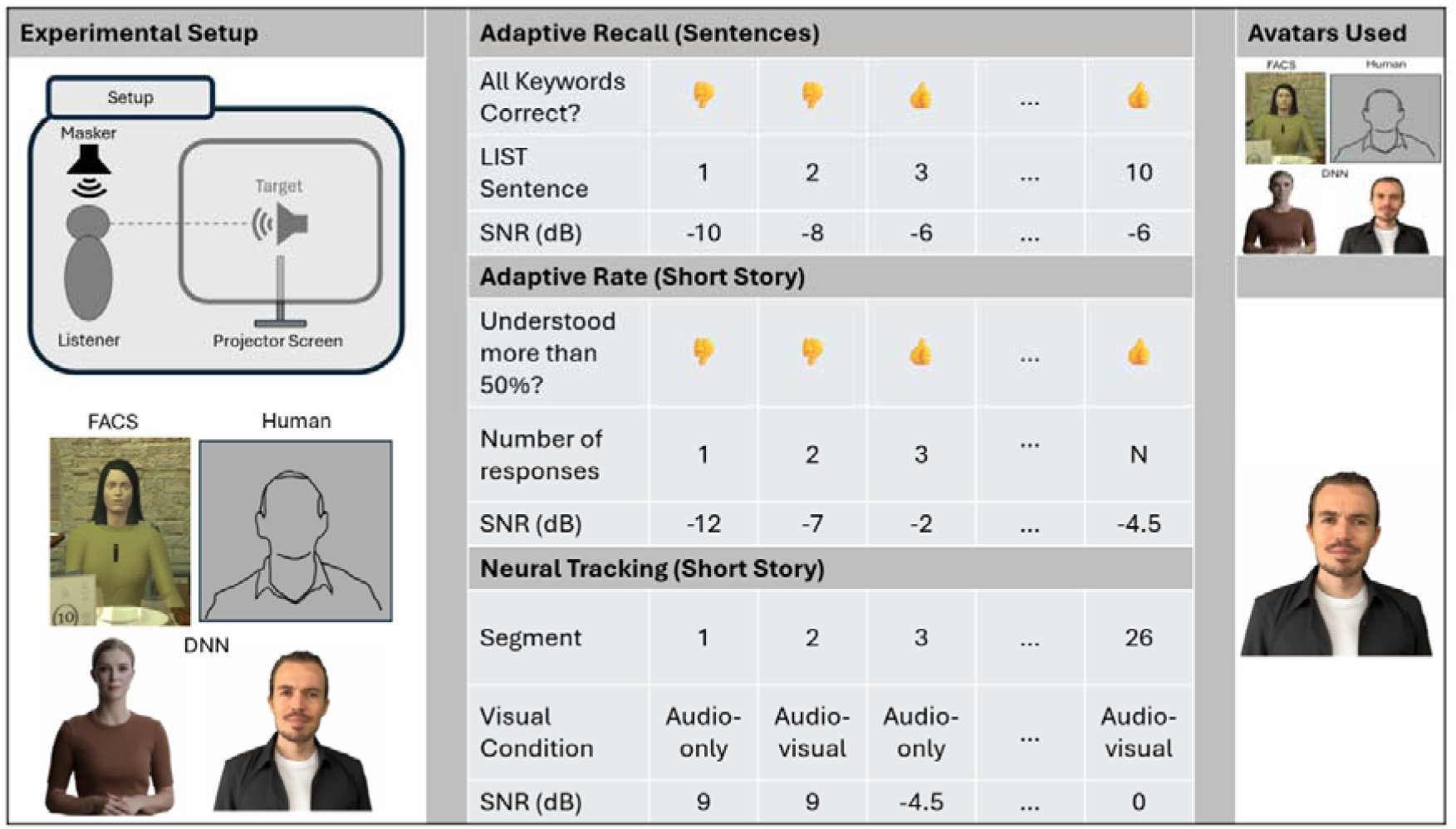
Experimental setup and procedures. Note that the avatars depicted in this figure are all computer generated and the video of the real human has been replaced with an outline. The top row of the first column presents an overview of the experiment setup. The bottom row of the first column illustrates four different speakers: an FACS avatar using the visual articulators, a human speaker, a deep neural network (DNN)-generated female speaker, and a DNN-generated male speaker. The top row of the milddle column depicts the first adaptive recall procedure using LIST sentences, utilizing all four speakers. The middle row of the second column showcases the adaptive rate procedure with LIST sentences, employing only the male DNN speaker. The bottom row of the second column demonstrates the neural tracking procedure utilizing a short story as stimuli, also using the male DNN speaker.

### Procedures

#### Behavioral

In this study, we utilized two behavioral measures to assess the audiovisual benefit on speech intelligibility. For the first behavioral measure, we employed an adaptive recall procedure (Devesse et al., 2018) using LIST sentences as the speech material. In the adaptive recall procedure, the signal-to-noise ratio (SNR) was adjusted in steps of 2 dB based on participants’ responses. Responses were scored on a sentence basis, marking a sentence as incorrect if any keywords were identified incorrectly. The initial SNR was set at −12 dB, and the first sentence was repeated until all keywords were correctly identified. Subsequent sentences were presented adaptively to estimate the SRT, converging toward a 50% understanding threshold. The SRT was calculated as the average SNR of the last five out of ten sentences, plus an eleventh imaginary sentence that would have been presented if the procedure had continued. Adjustments to the SNR were made by varying the level of multi-talker babble noise while keeping the target speaker level constant. Each participant engaged with two distinct lists of sentences for each speaker, categorized by visual condition into audio-only and audio-visual presentations. These visual conditions were presented sequentially in that specific order, resulting in a total of eight sentence lists per participant.

After the audiovisual condition for each speaker, participants evaluated the speaker by responding to three questions: “How realistic did you find the speaker to be?”, “How much benefit did the speaker provide for the task?”, and “How distracting was the environment around you while listening to the speaker?”. The response was a rating using a 7-point Lickert scale with 1 meaning “not at all” and 7 meaning “very much”. To familiarize participants with the experimental setup, we included a training phase using a separate list of sentences featuring both audio-only and audio-visual conditions presented via the FACS avatar.

For the second behavioral measure, we employed the adaptive rate procedure (Decruy et al., 2018), in which participants listened to two 1-minute segments of a continuous short story presented in background noise for each visual condition. They were instructed to inform the experimenter whether they understood greater than or less than 50% of the target stimulus at any given moment. Based on their responses, the noise level of the masker was adjusted in increments of 5 dB during the first three responses, followed by 2.5 dB for the remaining responses. This adaptive step approach was chosen to speed up convergence toward a speech reception threshold, given the constraints of limited experiment time and a finite number of responses. Participants had the flexibility to respond at any point during a segment and could submit multiple responses as they wished. To prevent participants from recalling the short story material for the subsequent electrophysiological experiment, we randomized the order of the story segments, ensuring no continuity between segments.

For the results of the adaptive rate procedure, to determine the speech reception threshold (SRT) levels for each visual condition, we fit a sigmoidal function to the response data for each participant using the optimize.curve_fit function from SciPy in Python (Virtanen et al., 2020). The response data consisted of binary values (0s and 1s), indicating whether participants understood less than or greater than 50% of the target stimulus at the different signal-to-noise ratios (SNRs). By fitting a sigmoid function to these binary responses, we were able to model the relationship between the SNR and the probability of understanding the speech stimulus. The estimated SRT level corresponds to the predicted midpoint of the fitted sigmoid function, representing the SNR at which participants had a 50% probability of correctly understanding the speech. We chose to fit a sigmoid to this response data instead of the averaging method used in the adaptive recall procedure because we wanted to use the sigmoid to predict the performance of each participant at each SNR level.

#### Electrophysiological

For the electrophysiological measure, we presented the 13 1-minute short story segments sequentially, each segment being presented twice, resulting in a total recording time of 26 minutes. The experimental variables included the visual condition and the signal-to-noise ratio (SNR) levels. The visual condition comprised two scenarios: audio-only (with no avatar on the screen) and audio-visual (with the avatar present). The SNR levels were set at −9, −4.5, 0, 4.5, and 9 dB. The 9 dB SNR condition was assigned a total of 5 segments per participant, while the other conditions were assigned 2 segments each. However, during data analysis, we opted to use 2 segments for each condition and trained a linear decoding model for all conditions (see below). The first two segments of the short story were always presented at 9 dB SNR, with the first segment being audio-only and the second being audio-visual, to help acclimate participants to the experiment and convey the main idea of the story. For the remaining segments, the conditions for the two experimental variables were selected randomly, ensuring a balanced representation across the experiment.

#### Neural Tracking

The preprocessing steps for this study were adapted from (Cooper et al., 2025). We utilized MATLAB R2022b and custom scripts (The MathWorks Inc., 2022) to preprocess the electrophysiological data. Initially, we high-pass filtered the data using a second-order zero-phase Butterworth filter with a 0.5 Hz cutoff frequency. The raw data were then epoched into 1-minute segments, resulting in a total of 26 segments, with 13 segments for each visual condition and 2 segments for each SNR condition, except for the 9 dB condition, which was trimmed down from 5 segments to 2 segments to match the other conditions. To reduce computational time for subsequent steps, we downsampled the data to a 256 Hz sampling rate using MATLAB’s built-in resample function. We employed a multichannel Wiener filter to remove eye blink artifacts from each EEG segment (Somers et al., 2018) and re-referenced the EEG channels to the average reference across channels. We then band-pass filtered the data between 0.5 and 15 Hz using a zero-phase Chebyshev type II filter, applying 80 dB attenuation at 10% outside the passband and maintaining a passband ripple of 1 dB.

We calculated the envelope of the speech material using a gammatone filterbank followed by a power law, as detailed in (Vanthornhout et al., 2018). The envelope was then downsampled to 256 Hz to match the sample rate of the EEG data. Subsequently, we filtered the envelopes between 0.5 and 15 Hz using the same band-pass filter applied in the EEG analysis. For both the EEG and the envelope, we concatenated the corresponding conditions and downsampled the signals to 64 Hz to reduce computation time for the neural tracking measures. Finally, we normalized the EEG and envelope using z-scoring per channel and per condition.

For our neural tracking measure, we used a participant-dependent linear decoder to reconstruct the speech envelope from the EEG. To train this decoder for each condition, we used ridge regression and 120 seconds of EEG and speech envelope data split using a fixed fold length of 30 seconds for K-fold cross-validation. The ridge parameter was set to the maximum value of the autocorrelation matrix of the EEG signal, which has been found to yield stable results in our lab. For the integration window, we chose 0 to 250 ms. We computed a Pearson correlation with the reconstructed envelope and the true envelope in each fold and averaged these correlations across folds to obtain reconstruction accuracy as a measure of neural tracking.

To estimate if there was a significant effect between reconstruction accuracy and SNR level, we computed a linear mixed effect model (Bates et al., 2015) in R (R Core Team, 2021) using the neural tracking data, according to the following equation:

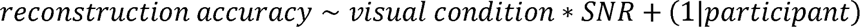

Where reconstruction accuracy defines the performance of the linear decoder, visual condition is the presence of a visual stimulus (audio-only and audio-visual) and participant is an individual participant. In this equation, we modeled the main effect of modality and SNR, their interaction effect (denoted by *), and modeled the participant as a random variable (denoted by (1|…)).

For all of our procedures we used non-parametric Wilcoxon tests from Scipy in Python (Virtanen et al., 2020) to perform pairwise comparisons for the assessment of differences between groups.

## Results

Behavioral testing consisted of both adaptive recall and adaptive rate procedures. For the analysis of the adaptive recall procedure, we found the audio-only conditions of the FACS avatar were not saved during the experiment for 2 participants; therefore, we excluded the data of these participants from the other conditions and speaker types. We kept these participants in the results of the other procedures as we do not directly compare the adaptive recall procedure results with any other procedure. The analysis of the adaptive recall procedure results (shown in Figure 2) revealed consistent audiovisual benefits across all speaker types: FACS, DNN, and Human. The magnitude of audiovisual benefit varied between speaker types, with the Human speaker providing the greatest benefit with an average SRT difference of −4.75 dB SNR. In contrast, the FACS and DNN speakers provided less benefit with average SRT differences of −1.55 dB SNR and −4.00 dB SNR, respectively. The SRT was different for each of the audio-only conditions because we used different sentences for each speaker type.

**Figure 2.**
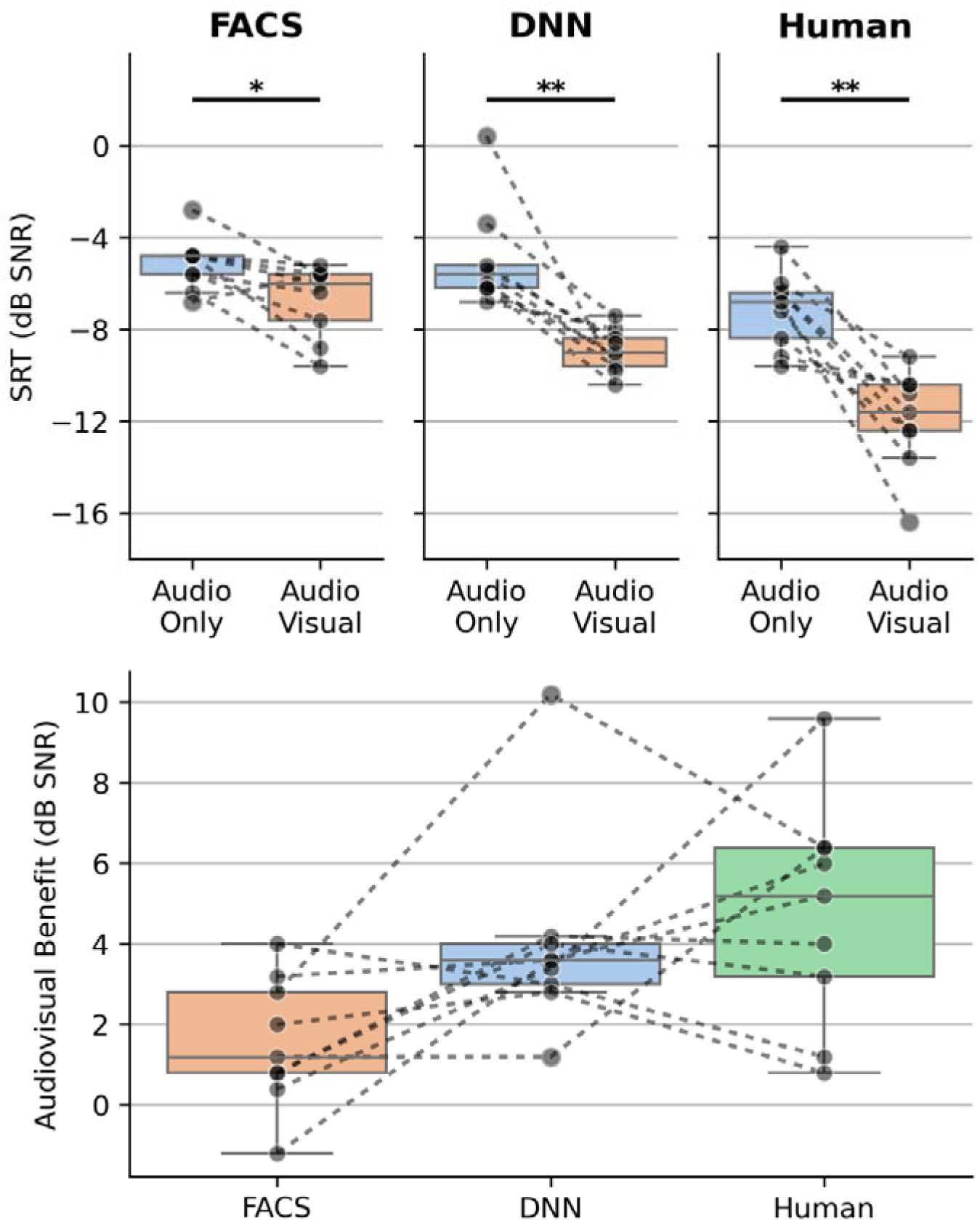
The top row depicts the speech reception thresholds (SRTs) obtained from the adaptive recall procedure for each participant across different speaker types and visual conditions. The bottom row illustrates the individual audiovisual benefit, calculated as the difference in SRTs between audiovisual and audio-only conditions for each speaker type. The DNN speaker type represents the average of the two speakers (male and female) across both panels. * denotes p-value < 0.05, ** denotes p-value < 0.01.

To assess differences in SRTs between visual conditions for each speaker type, we used Wilcoxon tests. From these tests, we found that there is a difference in SRTs between visual conditions for the FACS-based avatar (statistic= 40.5, non-adjusted p-value= 0.015625), the DNN-based avatar (statistic= 45.0, non-adjusted p-value=0.002), and the Human (statistic= 45.0, non-adjusted p-value=0.002).

To assess the audiovisual benefit, which is the difference between the SRTs in the audiovisual and audio-only conditions, we used Wilcoxon tests. We found that after comparing for multiple comparisons there were no differences in audiovisual benefit between each group (FACS-DNN: statistic=33.0, non-adjusted p-value=0.019, FACS-Human: statistic=40.0, non-adjusted p-value=0.019, DNN-Human: statistic=26.0, non-adjusted p-value=0.367).

Subjective ratings revealed differences in perceived realism and benefit across speaker types (see Figure 3). On average, the human speaker received the highest ratings for both realism (6.55) and benefit (6.27), followed by the DNN avatar (realism: 5.09; benefit: 5.18), while the FACS avatar received the lowest ratings (realism: 3.55; benefit: 3.91).

**Figure 3.**
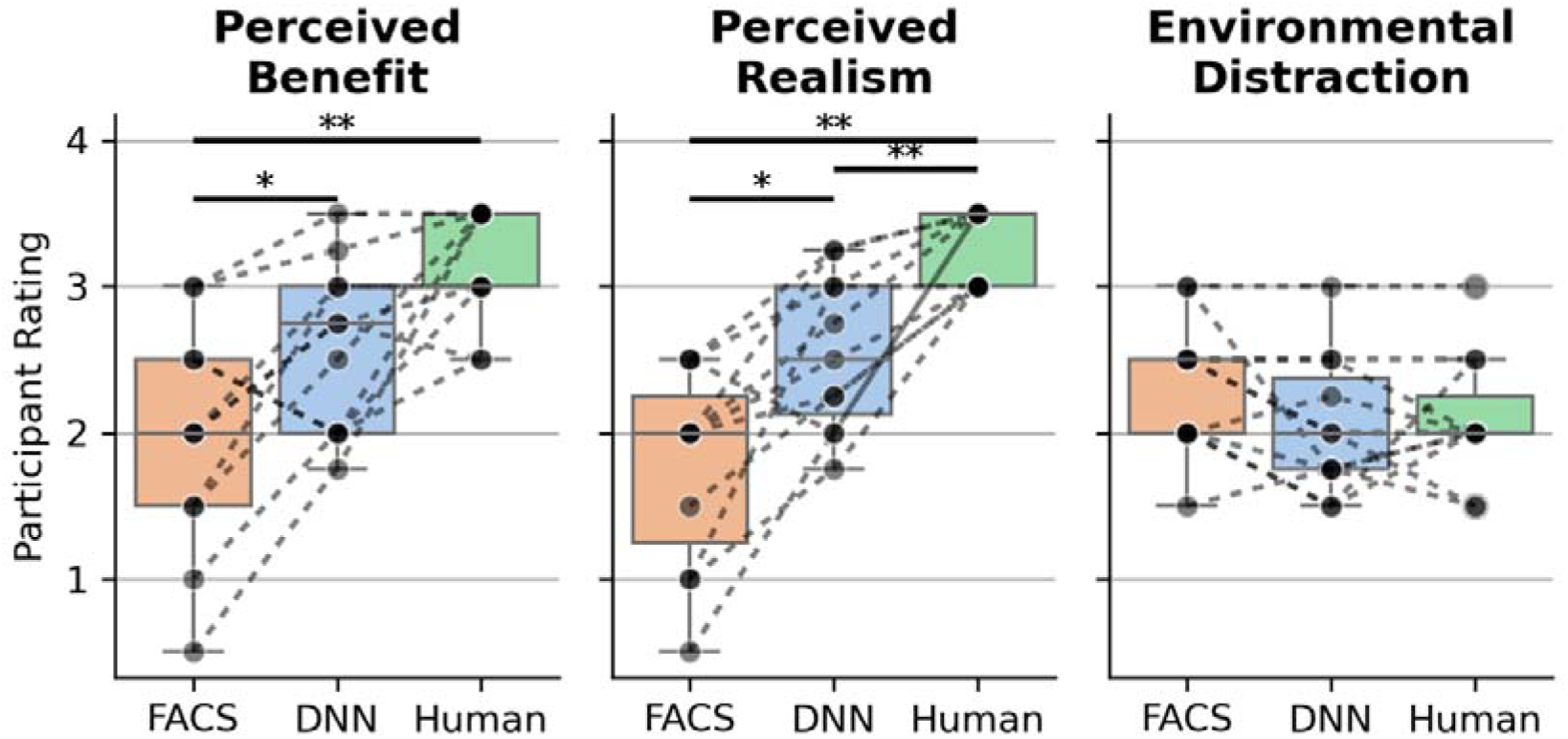
Subjective ratings across three dimensions (perceived realism, perceived benefit, and environmental distraction) following the audiovisual adaptive recall procedure for each speaker type. Ratings were collected on a 7-point Likert scale where participants evaluated: (1) the perceived realism of the speaker (1 = “not at all realistic,” 7 = “very realistic”), (2) the speaker’s effectiveness as a communication aid (1 = “not at all beneficial,” 7 = “very beneficial”), and (3) the level of environmental distraction during the task (1 = “not at all distracting,” 7 = “very distracting”). DNN speaker ratings represent averaged scores from both male and female DNN variants. * denotes p-value < 0.05, ** denotes p-value < 0.01, and *** denotes p-value < 0.001.

Pairwise comparisons using Wilcoxon tests revealed that each speaker type was different from each other in realism ratings (FACS-DNN: statistic=3.0, non-adjusted p-value=0.005, FACS-Human: statistic=0.0, non-adjusted p-value=0.001, DNN-Human: statistic=0.0, non-adjusted p-value=0.002). For the benefit ratings the test revealed a difference between FACS and Human and a difference between FACS and DNN (FACS-DNN: statistic=6.0, non-adjusted p-value=0.013, FACS-Human: statistic=0.0, non-adjusted p-value=0.002, DNN-Human: statistic=2.0, non-adjusted p-value=0.031). The perceived environmental distraction levels were comparable across all speaker types, with no differences observed (FACS-DNN: statistic=4.0, non-adjusted p-value=0.055, FACS-Human: statistic=6.0, non-adjusted p-value=0.25, DNN-Human: statistic=17.0, non-adjusted p-value=0.992).

In the second phase of the experiment, we examined both behavioral and neural responses to a short story delivered by a DNN avatar. The behavioral adaptive rate procedure demonstrated enhanced speech intelligibility in the audiovisual condition compared to audio-only presentation, with average SRTs improving from −7.17 dB SNR (audio-only) to −9.02 dB SNR (audiovisual), representing an audiovisual benefit of −1.85 dB SNR (see Figure 4). A Wilcoxon test revealed this difference (statistic = 0.0, p-value = 0.001). For the neural tracking results (shown in Figure 5), we computed the difference between the audiovisual and audio-only neural tracking values to obtain an audiovisual benefit neural tracking measure. We find that a majority of participants had an audiovisual benefit above zero only in the −9 dB SNR condition, which is near the audiovisual SRT levels of the behavioral results. We evaluated if the median of the reconstruction accuracy distribution at −9 dB was different than zero using a one-sided Wilcoxon test for each SNR condition and found that there is no difference after correcting for multiple comparisons (statistic = 53.0, non-adjusted p-value = 0.041).

**Figure 4.**
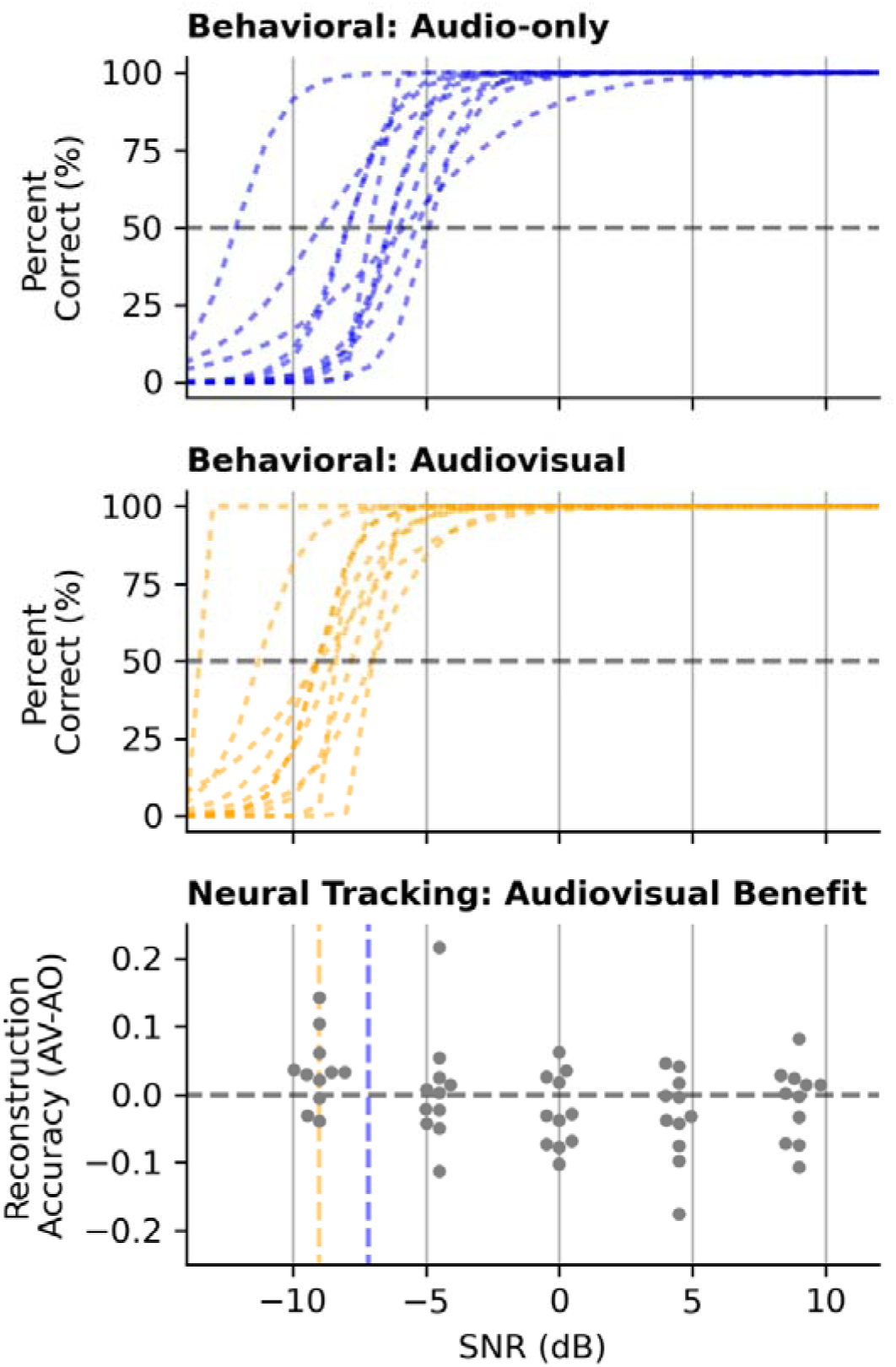
Comparing behavioral and neural tracking results. The top two rows show the sigmoidal fits from the adaptive rate procedure for each participant. The top row shows the fits for the audio-only condition and the middle row shows the fits for the audiovisual condition. The SRT level is marked with a dotted grey line at the 50 percent level. The bottom row shows the difference in reconstruction accuracies between the audiovisual and audio-only conditions for each SNR level and for each participant. The blue and orange dotted lines represent the average SRT level of all participants for the audio-only and audiovisual conditions, respectively.

**Figure 5.**
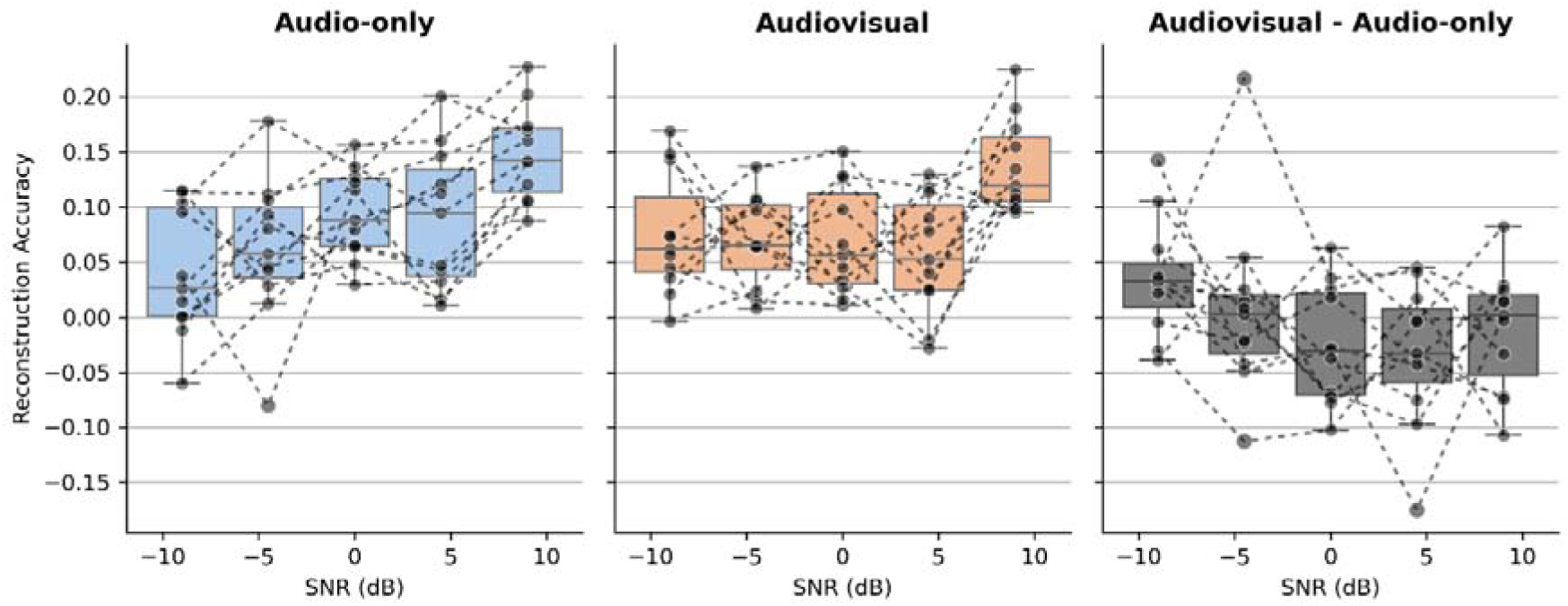
Comparison of neural tracking values. The left and middle columns show the reconstruction accuracies from the audio-only and audio-visual conditions, respectively. There are distinct neural tracking values for each SNR level and each participant. The right column shows the difference in reconstruction accuracy between the audiovisual and audio-only conditions for each SNR level and each participant. In each column the reconstruction accuracies associated with a participant are connected with a gray dotted line.

Using the linear mixed effects model on the neural tracking results, we examined the effects of SNR and visual condition on reconstruction accuracies. The model summary indicated that there was a main effect of SNR (Estimate = 0.00546, t-value = 5.229, p-value = < 0.001) and there was no main effect of visual condition (Estimate = −0.005365, t-value = −0.571, p-value = 0.569). Additionally, there was an interaction effect between the SNR and visual condition (Estimate = −0.00294, t = - 1.994, p-value = 0.049).

## Discussion

In this study, we employed three different procedures: a behavioral adaptive recall procedure, a behavioral adaptive rate procedure, and an electrophysiological neural tracking procedure.

In the adaptive recall procedure we evaluated the performance of three different speaker types (FACS avatar, DNN avatar, and Human) using one list of 10 sentences for each visual condition (audio-only and audiovisual). Our findings indicate that participants exhibited lower speech reception thresholds (SRTs) in the audiovisual condition compared to the audio-only condition. Notably, within the audiovisual condition, participants’ SRTs progressively declined from the FACS avatar to the DNN avatar, and ultimately to the human speaker. This trend, as corroborated from the results of the subjective ratings, suggests an increasing audiovisual benefit as the realism of the speaker type improves. We interpret this added realism as stemming from both the physical appearance of the speaker and the nuanced lip and facial muscle movements. Notably, the human speaker in the video recordings emphasized his mouth movements, which may have facilitated participants’ improved word identification. Additionally, non-verbal cues (e.g. arm and head movements) provided by the DNN avatar may have aided the participants more when compared to the absence of non-verbal cues provided by the FACS avatar.

Compared to the other speaker types, the DNN avatars offer greater flexibility for designing experiments that test audiovisual benefits with continuous stimuli. A primary advantage of using DNN models to generate virtual avatars is the ability to create audiovisual speech material from existing speech content. This approach can save significant amounts of money and recording time when producing a new audiovisual corpus based on an original audio-only dataset. Furthermore, we received comments from the participants that they did not realize the avatars used in our study were not real humans, which underscores the effectiveness of DNN avatar models. This inability to perceive the difference between real and synthetized videos of human talkers has been previously shown in an online Turing test (Varano et al., 2022).

For the behavioral adaptive rate procedure and the electrophysiological neural tracking procedure we found both show an audiovisual benefit when viewing the DNN male avatar reciting the short story. However, an audiovisual benefit observed in the neural tracking results only begins to emerge in the lowest SNR condition (−9 dB), which is around the behavioral SRTs. Holistically, these results showing a virtual human provides audiovisual benefit are in line with previous behavioral studies (Devesse et al., 2018; Shan et al., 2022; Varano et al., 2022; Yu et al., 2024) and neural tracking studies (Cooper et al., 2025), which all shown audiovisual benefit in the lower SNR ranges from −6 dB to −12 dB.

For the adaptive rate procedure we find that the DNN avatar yields an audiovisual benefit of 2 dB SNR, which is very similar to what has been found in previous studies (Devesse et al., 2018; Cooper et al., 2025). The similarity is a good validation of the DNN avatar as these previous results have been reported for the FACS avatar, which is structured based on anatomical principles to produce accurate visemes.

For the neural tracking procedure, we find that the median difference in reconstruction accuracies between audiovisual and audio-only are not different than zero for all the SNR conditions. However, the median difference in the −9 dB SNR condition may indicate a trend that lower SNR conditions would yield greater audiovisual benefit. An improvement to our experimental design would have been to include lower SNR levels than −9 dB to capture the increased audiovisual benefit; however, based on previous research with sentences (Cooper et al., 2025) and a short-story (Crosse et al., 2016) we hypothesized that −9 dB would already be a very difficult condition and yield the largest audiovisual benefit. Therefore, for future experiments, we plan to incorporate a −12 dB SNR condition.

From our analysis using a linear mixed-effects model, we found only an effect for the signal-to-noise ratio (SNR) levels and not for the visual conditions. We found that higher SNR levels are associated with increased correlation values. The absence of the effect of visual condition is expected since we observe that only from the −9 dB condition onward there is an emergence of a difference between the visual conditions. Additionally, the interaction between SNR level and visual condition suggests that when the visual condition is audiovisual, the effect of SNR is reduced. This makes sense considering our observations and previous research, which show that audiovisual cues can help listeners better understand speech, especially in situations with lower SNR levels (Crosse et al., 2016; Reisinger et al., 2025). A previous study, Cooper et al., 2025, found there was no interaction effect between SNR level and visual conditions, but this can be attributed to the inclusion of a silence condition. In this current study, noise is always present in each condition, which could account for the differences between studies. Overall, this highlights how audiovisual cues can improve the perception of speech and validates that the DNN avatar provides sufficient audiovisual benefit to improve speech perception.

We chose to use the envelope for our stimulus feature due to its importance in speech understanding (Shannon et al., 1995). Additionally, we expected the envelope to capture the slow fluctuations that would correlate with the mouth movements of the avatars enabling us to observe effects of audiovisual speech processing (Reisinger et al., 2025).

Although an improvement to our study would have been to utilize open-source models, we opted for a commercial model because of the realism achieved and its ease of use. We hope that commercial DNN models like the one used in this study will eventually be made available as open-source and free-to-use. Such developments would enable researchers to adjust various parameters of these models, allowing for further investigations into specific features that could enhance the audiovisual benefits observed in video recordings of humans. Progress has already been made in academic literature (Eskimez et al., 2018, 2020; Vougioukas et al., 2020; Eskimez et al., 2022). Additional open-source alternatives are currently available such as the Wav2Lip repository (Prajwal et al., 2020). This resource provides models and discriminators that generate realistic lip movements using a segment of audio and an existing video sequence. Further improvements could be achieved by employing stable-diffusion models on the video segments produced by Wav2Lip. A recent alternative, OmniHuman, that uses an end-to-end Diffusion Transformer-based framework may outperform all previous methods as the output videos provided from this method have realistic and temporally coherent lip movements and non-verbal gestures (Lin et al., 2025).

## Conclusion

In conclusion, our study demonstrates that from the adaptive recall procedure the audiovisual cues from virtual humans significantly enhance speech perception, with participants achieving lower speech reception thresholds (SRTs) in audiovisual conditions compared to audio-only conditions. This improvement was pronounced when using more realistic virtual humans, as evidenced by the progressive decline in SRTs from the FACS avatar to the DNN avatar, and ultimately to human speakers. Additionally, the behavioral adaptive rate procedure indicated a consistent audiovisual benefit of approximately 2 dB SNR for the DNN avatar, corroborating findings from prior studies. Neural tracking results further highlighted the effectiveness of audiovisual integration, showing a positive difference in reconstruction accuracies between the visual conditions only at the lowest SNR level (−9 dB), hence suggesting that participants leveraged visual cues to aid comprehension in noisy conditions. These findings reinforce the viability of DNN avatars as effective tools for studying audiovisual effects on speech perception.

